# The genetic legacy of archaic hominins in Central and Southeast Asia uncovers three distinct Denisovan populations

**DOI:** 10.64898/2026.05.06.723201

**Authors:** C. Antoine-Derouet, J. Adam Doucet, C. Leakhena Phoeung, C. Dorzhu, T. Hegay, E. Heyer, R. Chaix, C. Bon, F. Détroit, B. Toupance, R. Laurent

## Abstract

The sequencing of Neanderthal and Denisovan genomes has provided new insights into human evolution. Today, interactions between Neanderthals, Denisovans, and populations of European, East Asian, and Oceanian descent are well documented. However, neighboring regions such as Central and Southeast Asia remain understudied for archaic admixture despite their key geographic location and complex migration histories. To fill this gap, we investigate archaic ancestry in 16 populations from Central Asia and 14 from mainland Southeast Asia. Our results show that Neanderthal and Denisovan ancestry in these populations is of the same order of magnitude as in other Eurasian populations. However, although Denisovan ancestry accounts for less than 1% in mainland Asian populations, it originates from several admixture events involving different Denisovan populations. In particular, we find in Southeast Asia that Denisovan ancestry results from three distinct admixture events with three different Denisovan populations, highlighting the complexity of Denisovan contact with the ancestors of present-day Southeast Asian populations and providing new insights into the extensive geographic distribution of Denisovan populations.

## Introduction

Present-day Asian populations harbor a uniquely complex genetic legacy shaped by repeated waves of migration and admixture spanning the Late Pleistocene to recent historical periods (*1*–*6*). From the initial dispersal of anatomically modern humans out of Africa to later demographic events such as Neolithic expansions and historical trade networks including the Silk Road, Asia has played a central role in human population history. The continent also served as a major corridor for the peopling of Oceania and the Americas. These successive migrations and population interactions have profoundly shaped the genomic landscape of present-day Asian populations.

The sequences of archaic hominin genomes of Neanderthals (*7*) and Denisova from the Altai Mountains (*8*) provided direct evidence that admixture was not restricted to interactions among *Homo sapiens* populations, but also involved now-extinct hominins. These introgression events contributed to the genetic landscape of non-African Sapiens populations (*4, 8*–*15*). Early genomic studies documented widespread Neanderthal ancestry in European, East Asian, and Oceanian populations, estimating that Neanderthals contributed approximately 1-3% of the genomes of all non-Africans through a single admixture event (*10*–*12*). In contrast, Denisovan ancestry is limited to Asian and Oceanian populations. While East Asians harbor less than 1% Denisovan ancestry, populations from Oceania such as Papuans from Papua New Guinea and specific groups in Insular Southeast Asia like Agta from the Philippines, exhibit substantially higher levels, reaching at least 4% (*8, 12, 14*). This pronounced heterogeneity indicates that Denisovan admixture occurred through multiple, temporally and geographically distinct introgression events, likely involving genetically distinct Denisovan populations (*4, 12, 16*).

This genetic complexity is also mirrored by the Asian fossil record, which highlights the continent’s pivotal role in hominin evolution. Fossil remains attributed to *Homo erectus* from Trinil (Java, Indonesia) (*17*) and Zhoukoudian (China) (*18*), as well as more recent discoveries of *Homo floresiensis* (*19*), *Homo luzonensis* (*20*), and *Homo longi* (*21*), underscore the long-term persistence and diversity of hominin lineages in Asia. In recent years, additional debated taxa such as *Homo juluensis* have been proposed based on new analyses of fossil records from China (*22*), further emphasizing the taxonomic complexity of Late Pleistocene hominins in the region.

Importantly, molecular analyses have begun to clarify the evolutionary affinities of some of these enigmatic fossils. Proteins and mitochondrial DNA recovered from the Harbin skull (*23, 24*), initially described as *Homo longi*, and from the Penghu mandible (Taïwan) revealed that these individuals belonged to the Denisovan lineage (*23*–*25*). Together, these findings highlight both the evolutionary diversity of Denisovans and the difficulty of assigning Asian hominin fossils to taxonomic categories when molecular material is not available. When combined with genomic evidence for multiple Denisovan lineages (*4*), the fossil and molecular records jointly point to a complex and reticulated evolutionary history of hominins in Asia.

Despite recent advances, important gaps remain in our understanding of archaic introgression across Asia. Although a recent study demonstrated that *H. sapiens* Mainland Southeast Asian populations experienced admixture with multiple Denisovan populations (*26*), populations from Central Asia and Mainland Southeast Asia remain underrepresented in genomic studies of archaic ancestry. Central Asia shows a west-to-east genetic gradient. Western groups, particularly Indo-Iranian-speaking populations, are proximate to European and Iranian populations, while Eastern groups carry East-Steppe ancestry (*27, 28*) from admixture with westward-migrating nomadic populations during the 2nd–3rd century BCE (*3*). Mainland Southeast Asian populations exhibit genetic continuity with Neolithic individuals from the region (*1, 6, 29*) which themselves show admixture with Hoabinhian hunter-gatherers (*1, 6, 29*). This lack of representation is particularly striking given the geographic importance of these regions. Central Asia encompasses the Altai Mountains, where both Neanderthal and Denisovan remains were identified (*8, 13, 15, 30*), while Southeast Asia bridges East Asia and Oceania, two regions where Denisovan ancestry has been widely explored and shown to follow complex patterns. Moreover, both regions functioned as major corridors for human dispersal across Eurasia and into Sahul. Addressing this gap is therefore essential for reconstructing the spatial and temporal dynamics of archaic admixtures and for understanding its contribution to present-day Asian genetic diversity.

To address this gap we investigate the admixture between Neanderthal, Denisova and the ancestors of Central and mainland Southeast Asian populations. We leverage high coverage genomes from the Altai Neanderthal (*15*), Vindija Neanderthal (*11*) and Denisova 3 (*8*), alongside genetic data from Asian populations from the 1000 Genomes Project (*31*) and genome-wide data from Central, Northern and Southeast Asian populations generated by our group (*32, 33*).

## Results

### Population structure

We analysed genomic data from 427 individuals belonging to 16 populations from Central Asia (Fig. 1A-1B, Table S1) and 581 individuals from 14 populations sampled in Laos and Cambodia (*28, 32*–*35*) (Fig. 1A, 1C).

**Fig. 1.**
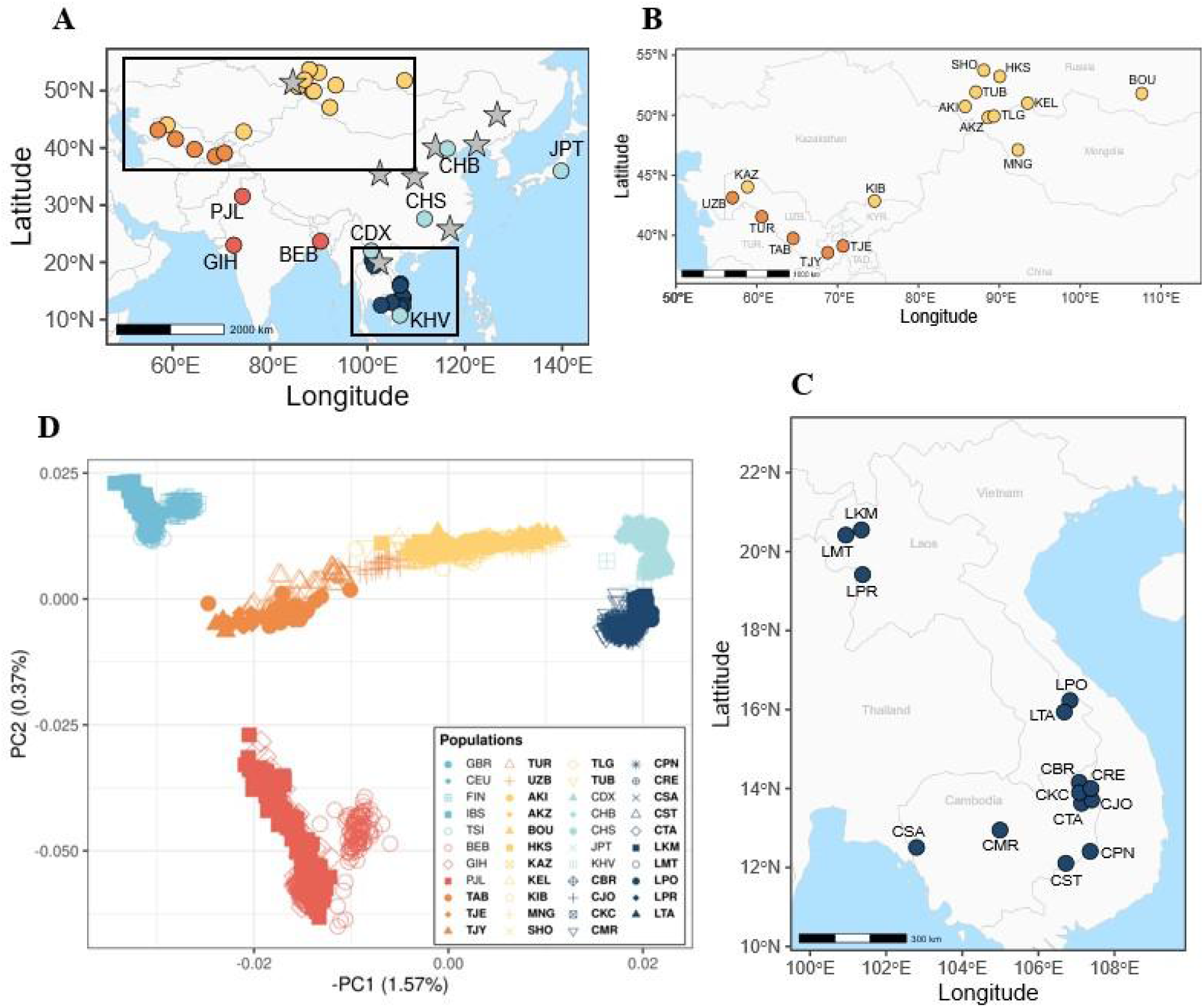
Populations of this study. **A**. Map showing populations from the 1000 Genomes database. Grey stars represent Denisovan fossils and fossils suspected to belong to Denisovans. **B**. and **C**. Map focusing on populations of this study. **D**. PCA on populations from the 1000 Genomes database and this study. Mid-blue, red, orange, yellow, light blue and dark blue points respectively represent European, South Asian, Central Asian, Northern Asian, East Asian and Southeast Asian populations. Population names of this study are in bold, and the full names are available in Table S1.

Principal component analysis (Fig. 1D), computed on contemporary human populations, reveals an East-West differentiation along the first principal component and a North-South differentiation along the second. Central and Northern Asian populations form a cline along a gradient from East Asian to European populations. The distribution of this genetic pattern mirrors that of linguistics, as previously described in (*2, 27, 28, 36*). Mainland Southeast Asian populations form a separate cluster, distinct from East Asians, with the exception of Kinh Vietnamese individuals, clustering instead with East Asian populations. Combining PCA results with geographic information, we classified populations into five groups: South, Central, Northern, East and Southeast Asia. Based on PCA results, we assigned KAZ and KIB to Northern Asian populations and KHV from the 1000 Genomes Project to East Asian populations (Table S1).

### Genome-wide Neanderthal and Denisovan ancestry

To estimate the extent of archaic ancestry in Central and Southeast Asian populations, we computed D-statistics (*37*) and f4-ratio statistics (*37*). We found that Neanderthal ancestry is homogeneous across Asian populations (mean = 2.77%, sd = 0.10), although it is slightly reduced in Central and South Asian groups (mean = 2.54%, sd = 0.04) (Fig. 2A). In contrast, Denisovan ancestry is highly heterogeneous across Asia (Fig. 2B). No detectable Denisovan ancestry is observed in Central or South Asian populations, whereas East, Northern, and Southeast Asian populations inherited less than 1% (mean = 0.62%, sd = 0.09) of their genome from Denisovans (Fig. 2B).

**Fig. 2.**
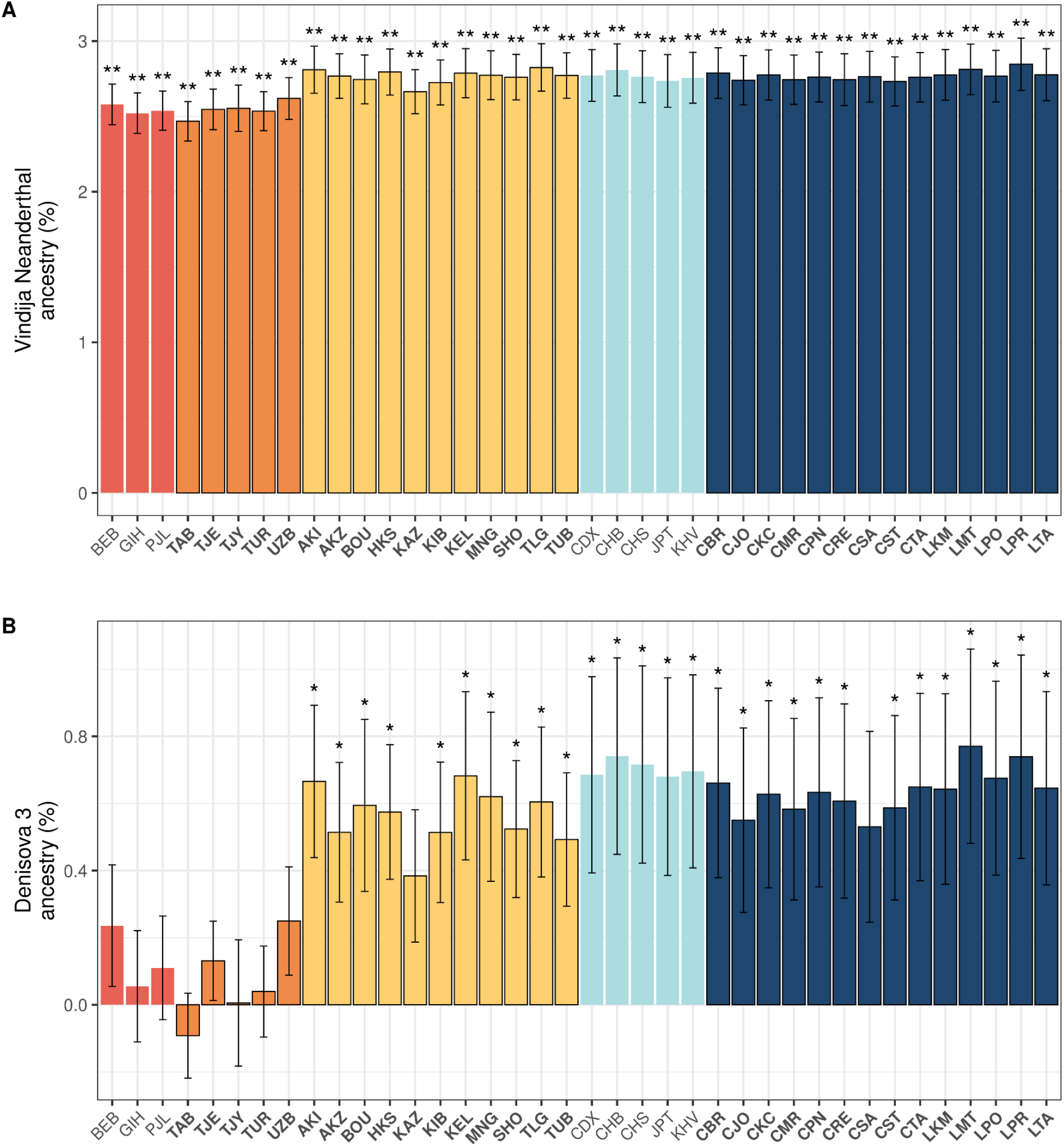
Archaic ancestry of Asian populations. **A**. Levels of Neanderthal ancestry using Vindija Neanderthal. **B**. Levels of Denisovan ancestry estimated using Denisova 3 genome. Error bars represent 2 standard deviations around the point estimate, computed using a weighted-block jackknife procedure. Population names of this study are in bold. A p-value lower than 0,01 is indicated by ** and lower than 0,05 by *.

### Multiple Denisovan introgression events revealed by haplotypes

To resolve the structure of Denisovan ancestry, we identified introgressed haplotypes combining two complementary approaches: Sprime (*4*), which detects introgression without requiring archaic references, and a conditional random field (CRF) approach (*10*), which incorporates Neanderthal and Denisovan genomes as references.

For all populations, we computed the match rate of each segment identified by Sprime as the proportion of sites matching the genome of Neanderthal or Denisova relative to the total number of sites in the segment. The distribution of these match rates to the Vindija Neanderthal genome shows a single peak at ∼0.9 (Fig. 3, S3), consistent with previous reports and supporting a single introgression event from a Neanderthal population into the ancestors of all non-African populations (*4, 12*). In contrast, Denisovan-derived segments displayed more complex patterns. A peak at a match rate of ∼0.6 is present in nearly all Asian populations, suggesting a widespread contribution from a Denisovan lineage moderately distinct from the Denisova 3 individual. A second peak at a match rate of ∼0.8 is detected in Northern, East and Southeast Asians, reflecting introgression from a Denisovan population more closely related to Denisova 3. Finally, several Southeast Asian groups (e.g., CTA, CJO, CST, LPO; Fig. S3) exhibit a third peak at ∼0.35, consistent with introgression from an additional deeply diverged Denisovan lineage (Fig. 3, S3). To confirm these observations, we combined haplotypes identified by Sprime with those identified by the conditional random field (CRF) approach developed by Sankararaman et al. (*10*) which detects archaic introgressed haplotypes using an archaic reference genome. We restricted further analyses to segments jointly identified by both methods to limit the number of false positive haplotypes (*12, 16*).

**Fig. 3.**
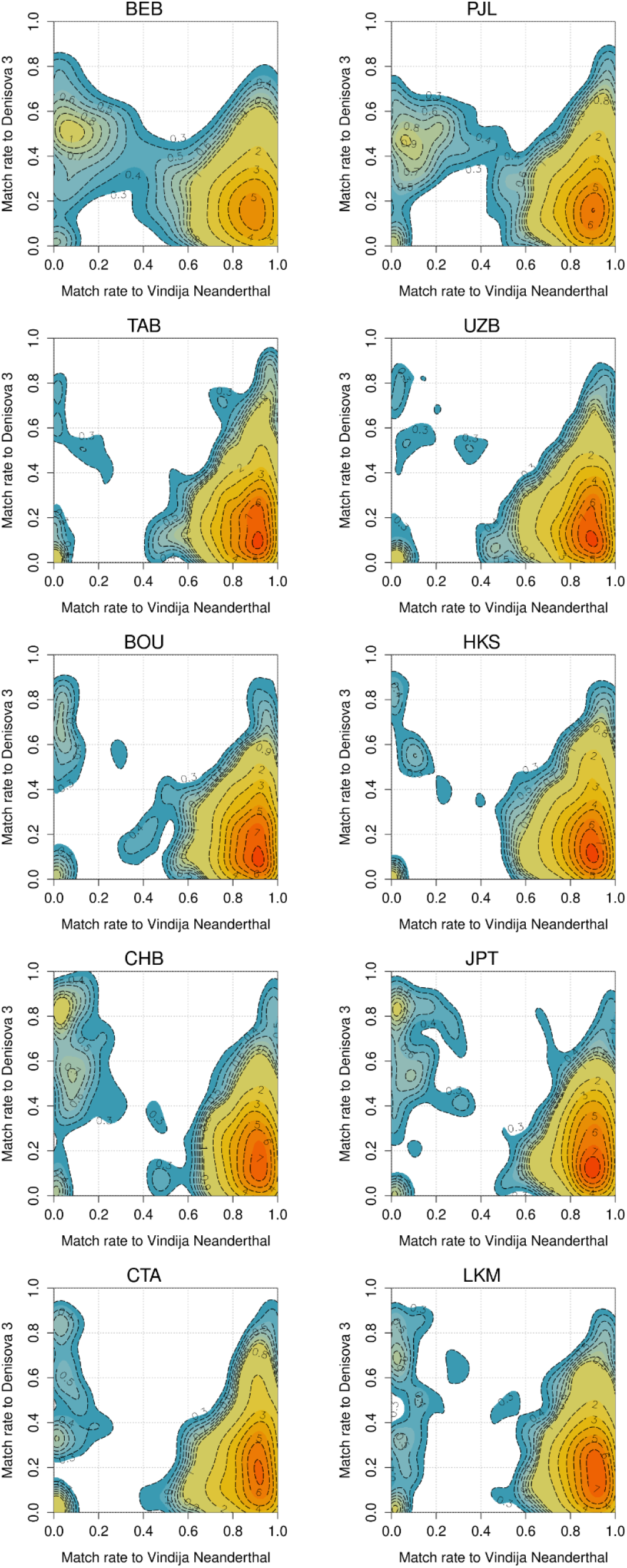
Contour density plots of match rates of introgressed segments to the Vindija Neanderthal and Denisova 3 genomes. Match rate is defined as the proportion of putative archaic alleles matching a given archaic genome, excluding masked sites. Only Sprime haplotypes with at least 30 unmasked sites in the Vindija Neanderthal and Denisova 3 genomes are included in the match rate calculations. Contour labels represent density levels. Contour lines are shown for multiples of 1 (solid lines) and multiples of 0.1 between 0.3 and 0.9(dashed lines). Colors were added to improve interpretability.

We first analysed the distribution of Neanderthal and Denisovan haplotypes across populations (Fig. S4) and then, to maximize the number of haplotypes available for further analyses, we merged haplotypes within each of the 5 population groups (Fig. 1, Table S1). Neanderthal match-rate distributions are unimodal, whereas Denisovan distributions are multi-modal (Fig. 4). In South, Central and East Asia, Denisovan match rates are bimodal with peaks at ∼99.2% and ∼98% (97.5% in South Asia), hereafter called Pulse1 and Pulse2, respectively (Table S2, S4, S5). These patterns suggest that Denisovan ancestry in these populations results from two different admixture events involving two Denisovan groups genetically differentiated from Denisova 3.

**Fig. 4.**
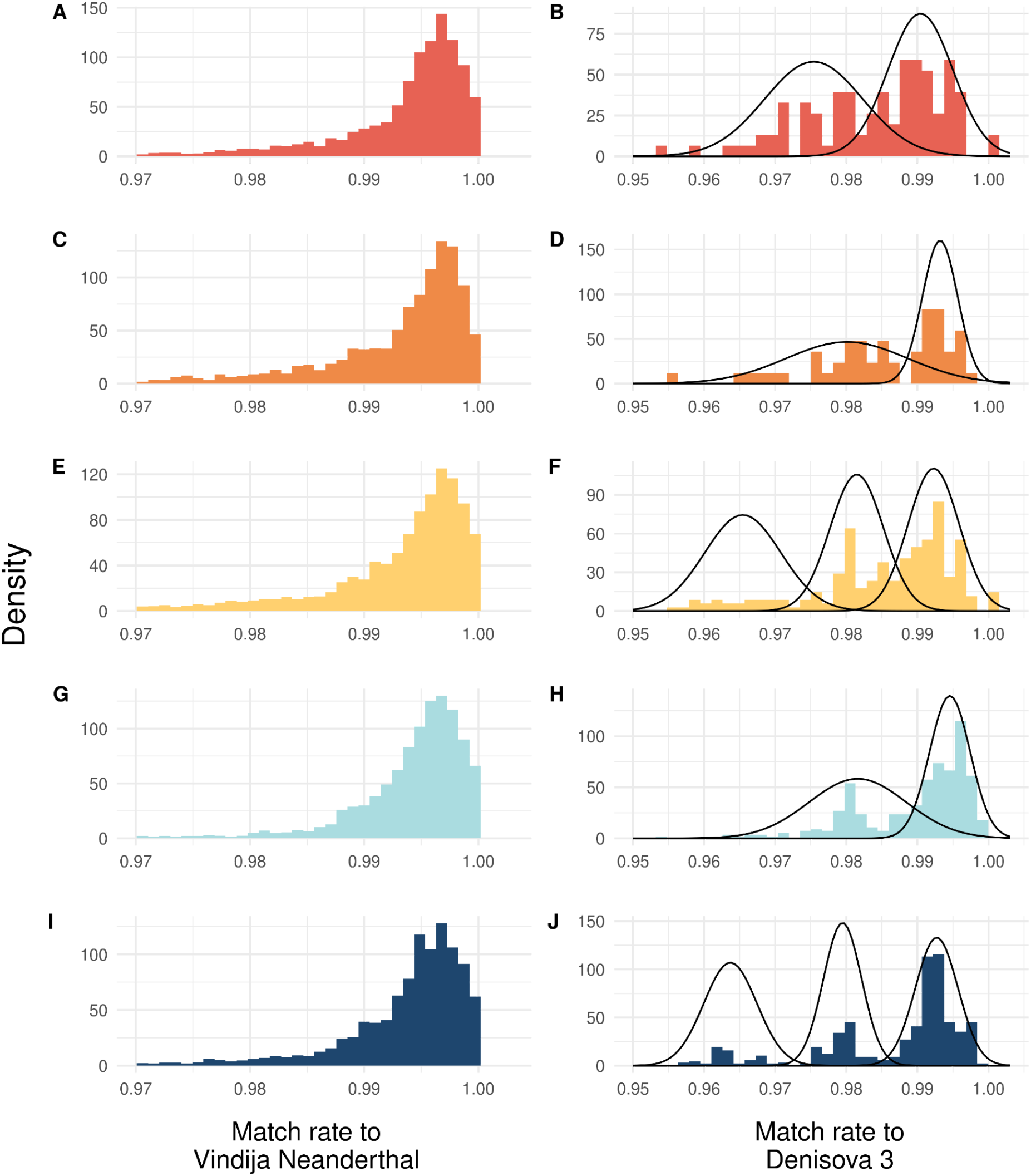
Match rate distributions of introgressed haplotypes identified by both Sprime and CRF. Gaussian mixture models were fitted to the distributions. Left (A, C, E, G, I): Vindija Neanderthal; right (B, D, F, H, J): Denisova 3. Solid curves represent fitted Gaussian mixture models

Signals associated with these two components are also observed in Northern and Southeast Asia populations. In addition, an extra peak at ∼96.4% (hereafter called Pulse3) (Fig. 4, Table S2, S5) indicates introgression from a more deeply diverged Denisovan population, which also contributed to the ancestry of Northern and Southeast Asian populations.

In South and East Asia, haplotypes in Pulse1 are longer than those in Pulse2, as previously reported (Fig. 5, Table S4) (*12*). However, in Southeast Asia, the pattern is reversed, with Pulse1 segments shorter than Pulse2 segments (Wilcoxon test, p < 0.01). Moreover, Neanderthal haplotypes are generally longer than Denisovan haplotypes (Table S6).

**Fig. 5.**
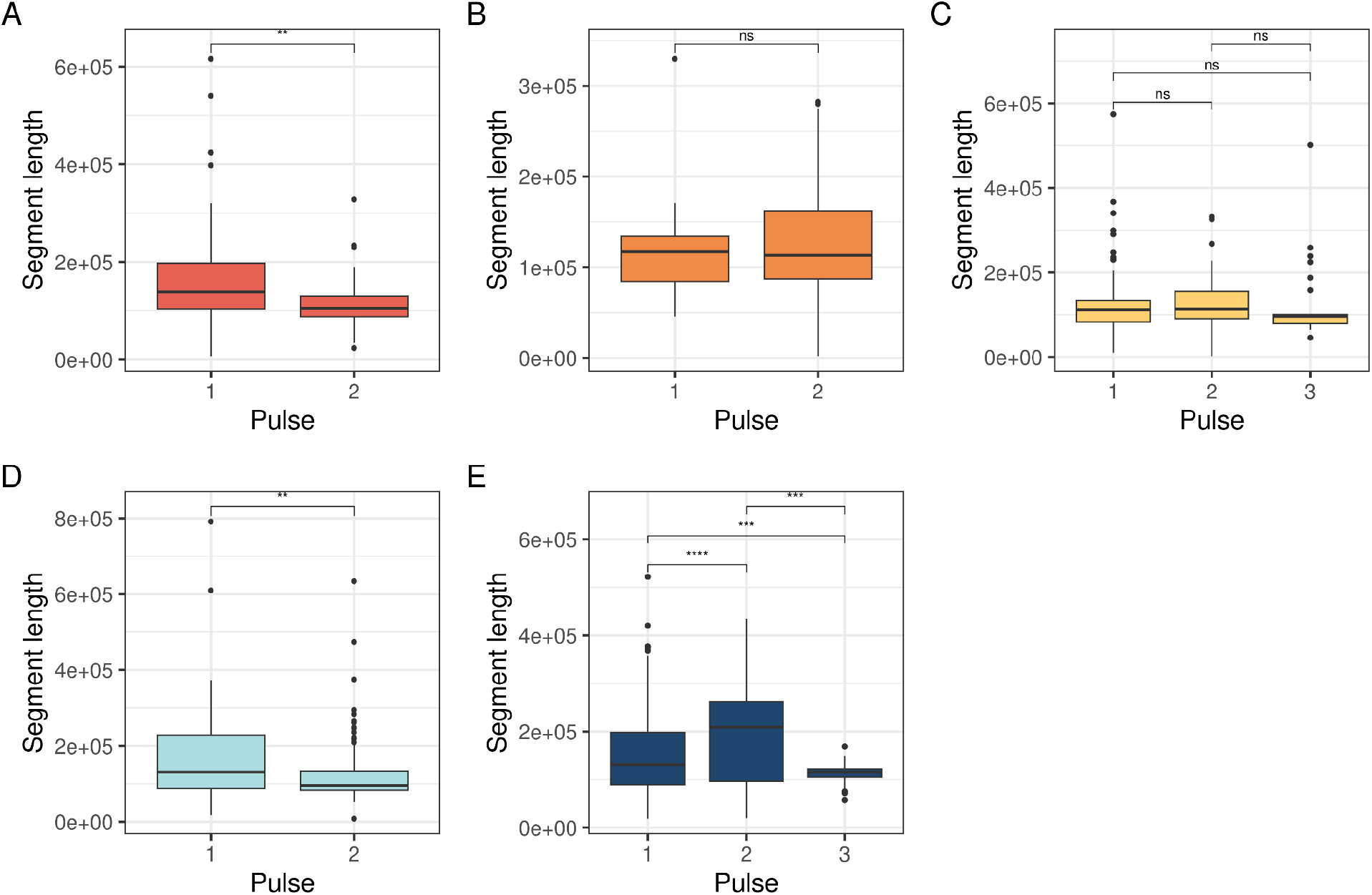
Haplotype length distribution across admixture pulses. Haplotype lengths assigned to each pulse were compared using Wilcoxon rank-sum tests. Haplotypes in Pulse1 have the highest match rates, those in Pulse2 have moderate match rates and those in Pulse3 have the lowest match rates. A: South Asia, B: Central Asia, C: Northern Asia, D: East Asia and E: Southeast Asia.

Overlap analyses reveal that Pulse1 segments, as well as Pulse2 segments, broadly overlap across most Asian regions, particularly for the Pulse1, consistent with a single admixture event with a common Denisovan source population (Fig. S7). The overlap is reduced in South Asia, especially for Pulse2. In contrast, Pulse3 shows weak overlap between Northern and Southeast Asia, relative to other pulses, casting doubt on a single common origin (Fig. S7).

### Phenotypic consequences of archaic introgressions

It has previously been shown that archaic variants inherited from Neanderthals and Denisovans can influence phenotypes in modern human populations. We therefore examined whether the archaic segments we identified contain genes reported in previous studies (*4, 12, 38, 39*). Genes located in Neanderthal haplotypes were mainly involved in immune-related functions, such as *CXCR6* and *IL10RA*, metabolic processes, such as PASK and *OSBPL10*, and neuronal development, such as *TENM3* and *MCPH1* (*4, 12, 40*–*42*). We also found that Neanderthal haplotypes in East and Southeast Asia overlap the *HS3ST3A1* gene, involved in tooth morphology and previously reported as introgressed from Neanderthals (*43*). Denisovan haplotypes identified in East and Southeast Asia include immune-related genes, such as *BANK1* (*12*).

### Adaptive introgression

Admixture with Neanderthals and Denisovans introduced adaptive variants that contributed to human adaptation to novel environments (*12, 38, 44*). We assembled a list of genes located at adaptive introgression sites that were described in literature and searched for genomic regions harboring both i) introgressed segments and ii) signatures of positive selection based on iHS. We detected four loci under positive selection within Neanderthal-introgressed segments (Table S7). Signals of positive selection were identified at CNTN5 in Northern and Southeast Asian populations. This gene is implicated in brain neurodevelopment (*43, 44*), and signatures of selection have previously been reported in different Philippine and Southwestern Chinese populations (*12, 47*) (Fig. 6A). TENM3, also involved in brain neurodevelopment (*48*), was identified as an adaptive introgressed locus in NA, CA and SEA. Previous studies have reported signals of selection at this locus in Philippine and Polynesian populations (*12*). Lastly, signatures of positive selection were identified within MCPH1 and TBC1D1 introgressed segments. MCPH1 is involved in cerebral cortex development (*45*), whereas TBC1D1 is associated with metabolic functions (*50*).

**Fig.6.**
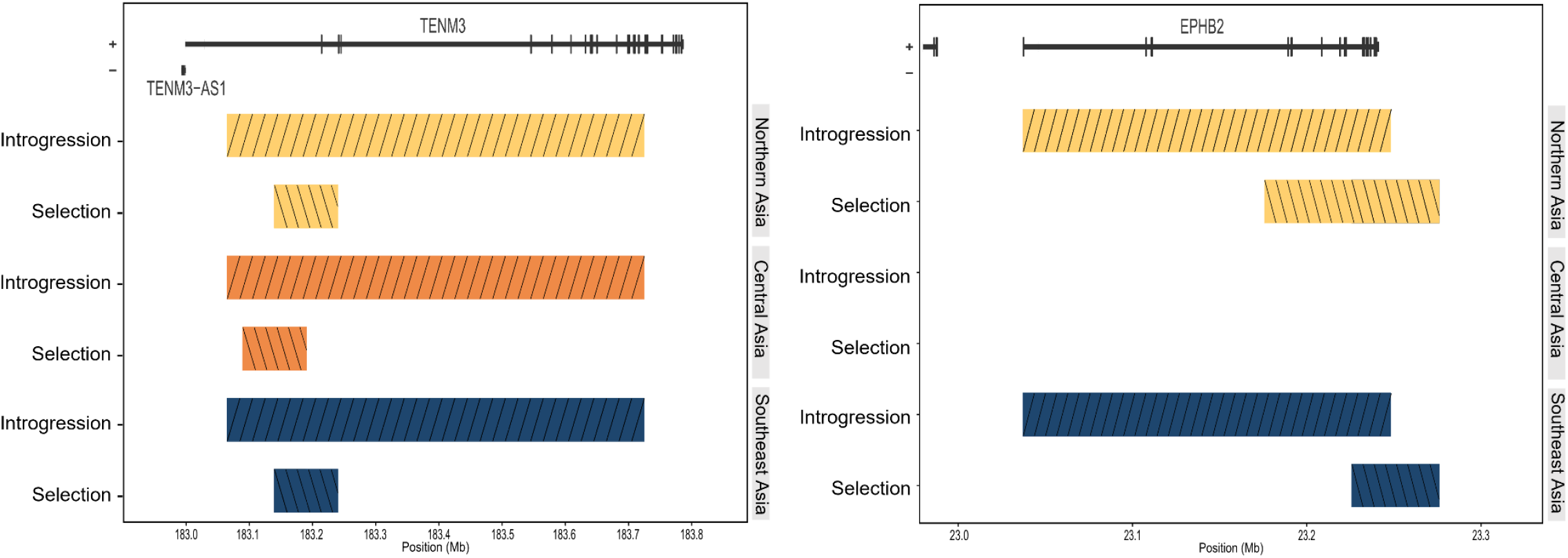
Examples of adaptive introgression regions from Neanderthal and Denisova. Colored rectangles represent the genomic intervals where introgression or selection was observed in each géographic region. The positions of overlapping genes are represented at the top. **(A)** Introgressed segments from Neanderthal. **(B)** Introgressed segments from Denisova.

We also identified three genomic regions under positive selection within Denisovan introgressed segments, encompassing BANK1, EPHB2 and FLG/HRNR (Table S7). BANK1 and EPHB2 show signatures of positive selection in SEA populations for the former, and in SEA and NA populations for the latter (Fig. 6B). Both genes are involved in immune functions and have previously been reported as adaptive introgression loci in Philippine and Taiwanese populations, respectively (*51, 52*). The last loci encompasses FLG and HRNR and is found under selection in SEA populations. These genes are involved in skin barrier function, keratinocyte differentiation, and hair shape, and have been described as part of a population-specific sweep in Han Chinese populations (*53, 54*).

## Discussion

Our analyses provide new insights into the genetic history of Asian populations along two major routes of human peopling of Eurasia, through Central and Southeast Asia. Neanderthal ancestry is estimated at approximately 2.7% (mean ≈ 2.72, sd ≈ 0.099) across Central, Northern, and Southeast Asia, in agreement with levels reported in other Eurasian populations (*11*). By contrast, Denisovan ancestry occurs at low levels in Northern and Southeast Asia, resembling patterns observed in East Asians (*4*) but exhibiting a more complex structure.

Two Denisovan admixture pulses, with match rates of 99.2% and 98% (Pulse1 and Pulse2), are widespread across Asia (Fig. 3, 4, S3, S4, Table S2, S4, S5). Pulse1 haplotypes, closely related to Denisova 3, are detected across all regions. One possible explanation, consistent with previous hypotheses, is that this admixture occurred in East or Northern Asia, among the common ancestors of Northern and East Asians, and that these Denisovan-derived haplotypes later entered Southeast Asian populations during Neolithic expansions (*1, 4, 6, 29*), as suggested by the overlap of shared haplotypes. In South Asia, this two-pulse signal is observed only in the Bengali (BEB) population, a Northeastern of South Asia group with genetic affinity and recent admixture with East Asian populations, which may have introduced these haplotypes into the Bengali gene pool (*4, 55*).

Haplotypes showing a moderate match rate (around 0.6 as inferred from Sprime) to Denisova 3 were found, not only in Mainland Asian populations, but also in Oceanians (*12, 16*). However, the genomic distribution of Denisovan segments in ancestral East Asians is not consistent with that in present-day Oceanians, suggesting that this ∼0.6 component does not reflect a single shared admixture event between Denisova and the ancestors of present-day Asians and Oceanians (*3*). The origin of this Denisovan ancestry in Mainland Southeast Asians therefore remains uncertain. Gene flow from Insular Southeast Asia or Oceania is one possible source, although current evidence does not strongly support this hypothesis (*56*). This underlines the need to better characterize the genetic relationships among Oceanian, Island Southeast Asian, and Mainland Asian populations. In South Asia, Denisovan haplotypes, particularly those in Pulse2, weakly overlap with those from other Asian groups (Fig. S7), suggesting possible inheritance from a distinct Denisovan population (*57*).

A third, more divergent pulse with a 96.4% CRF match rate to Denisova 3 appears geographically restricted to Northern and Southeast Asia (Fig. 4, Table S5). Denisovan ancestry in Southeast Asians has previously been proposed to result from three different contacts with three different Denisovan populations (*25*), and our results provide additional support for this hypothesis. Evidence for a three-pulse signal in Northern Asia remains limited, due to the small number of introgressed segments (Table S5) and its absence in Sprime-based analyses (Fig. 3, S3). If confirmed, this pattern would imply that multiple admixture events also occurred in Northern Asia, involving distinct Denisovan sources. Highly divergent Denisovan haplotypes, strongly distinct from Denisova 3, have also been reported in Oceanian (*12, 16*) and Island Southeast Asian populations (*14*). Such divergent haplotypes likely reflect admixture with a highly distinct Denisovan population, possibly in Mainland Southeast Asia or Insular Southeast Asia, later reintroduced into Mainland Southeast Asian populations through more recent interactions. Alternatively, different Denisovan populations may have been encountered independently by the ancestors of Southeast Asians and by Insular Southeast Asians and Oceanians.

Haplotype length can provide reliable insights to infer the relative timing of admixture events. In South and in East Asia, haplotypes most closely matching the Denisova 3 genome (Pulse1) tend to be longer than moderately matching segments (Fig. 5, Table S4), suggesting that admixture with the Denisovan population closely related to Denisova 3 is the most recent, in agreement with previous studies (*12*). In contrast, in Southeast Asia, the moderately related Denisovan pulse (Pulse2) shows the longest haplotypes, whereas the more divergent pulse (Pulse3) shows the shortest (Fig. 5, Table S5). In Central and Northern Asia, haplotype length does not differ significantly among pulses (Fig. 5, Table S4, S5). Unexpectedly Denisovan haplotypes were on average shorter than Neanderthal haplotypes (Table S6), even though Denisovan admixture has been estimated to be ∼10% more recent (*10*), with Neanderthal gene flow dated to 50–60 kya (*59, 60*).

Overall, these results reveal the complexity of human migrations and archaic admixture in Asia, where multiple scenarios may explain the observed Denisovan ancestry patterns. To reconstruct this intricate history, future demographic models must integrate genetic data from Mainland Asia, Island Southeast Asia, and Oceania. Our results also inform hypotheses about Denisovan geographic distribution. The presence of low-match Denisovan pulses in Papuan (*12, 16*) and Southeast Asian populations (both Mainland and Insular) supports encounters with Denisovans present in Southeast Asia. Although sequencing of additional Denisovan genomes has expanded nuclear DNA datasets, all currently published genomes originate from the Altai Mountains, and no genome has yet been recovered from southern regions where admixture with ancestors of Southeast Asians likely occurred, possibly due to poor preservation in tropical contexts. The discovery of a Denisovan-affiliated molar (*61*) further supports Denisovan presence in these areas.

Recent mitochondrial and proteomic evidence suggests a broader Denisovan range extending to northern China, the Tibetan Plateau, and Taiwan (*23*–*25, 62*). However, the absence of nuclear genomes from Southeast Asia and Oceania remains a key limitation for refining models of Denisovan introgression and dispersal. Progress will require additional ancient DNA data from lower latitudes, where possible, and the testing of explicit demographic models that jointly include Asian and Oceanian populations to distinguish among alternative scenarios of admixture, structure, and expansion. This will demand considerable effort given the climatic and preservation challenges in Southeast Asia.

In this study, we provide new insights into the genetic history and admixture processes of Asian populations that have been underrepresented in genetic studies, revealing evidence for three distinct Denisovan sources in Southeast Asian genomes. These findings emphasize the pivotal role of this region in past interactions between modern humans and archaic hominins.

## Materials and Methods

### Experimental design

This study aimed to quantify and characterize Neanderthal and Denisovan ancestry in Central and Southeast Asian populations, and to refine our understanding of the demographic history of archaic introgression in Mainland Asia.

We first estimated genome-wide levels of archaic ancestry using D-statistics and f4-ratio statistics (*37*).

To characterize the structure and timing of Denisovan introgression, we focused on introgressed haplotypes. Putative archaic haplotypes were initially identified using Sprime, and we computed their genetic affinity to the Denisova 3 genome (*8*). To reduce false positives and retain high-confidence archaic segments, we intersected Sprime-inferred haplotypes with those detected using a Conditional Random Field (CRF)-based approach (*10, 12, 16*). For the retained segments, we recalculated their genetic proximity to Denisova 3 (match rate).

To determine the number of distinct Denisovan introgression events experienced by the ancestors of Mainland Asian populations, we modeled the distribution of Denisovan match rates using Gaussian mixture models (GMMs) and selected the optimal number of components using the Bayesian Information Criterion (BIC). Each inferred component was interpreted as a distinct Denisovan pulse of admixture.

To infer the relative timing of introgression events, we compared the length distributions of haplotypes assigned to each pulse using Wilcoxon rank-sum tests, under the expectation that older introgression events generate shorter segments due to recombination.

Finally, we screened introgressed Neanderthal and Denisovan haplotypes for signatures of adaptive introgression.

### Populations and datasets

We analysed data from 30 populations (1008 individuals) in Central, Northern and Southeast Asia, from two former projects (*28, 32*–*35*) led by Evelyne Heyer and Raphaëlle Chaix from the Eco-Anthropology lab. A total of 912 individuals from 28 populations were genotyped using Omni1 and/or Omni2.5 SNP arrays. The remaining 96 individuals, representing 6 populations, are whole-genome sequenced samples (30 X coverage) (with 4 populations containing a mix of genotyped and whole-genome sequenced individuals, Table S1).

The Central Asia dataset includes genomic data from 427 individuals belonging to 16 populations spanning a broad definition of Central Asia, generated using whole-genome sequencing and Illumina Omni1 and Omni2.5 genotyping arrays. Based on linguistic affiliations, these populations can be broadly divided into Indo-Iranian speakers, who show greater genetic affinity with European populations, and Turko-Mongol speakers, who are more closely related to East Asian populations (*27, 28, 36*). TheTurkmen population represents a notable exception, as it speaks a Turko-Mongol language but clusters genetically with Indo-Iranian-speaking groups (*27, 28, 36*).

In Mainland Southeast Asia, the dataset includes 581 individuals from 14 populations sampled in Laos and Cambodia. These populations are predominantly Austroasiatic speakers, with the exception of the Jarai, who speak an Austronesian language (*32*). They are thought to derive ancestry from both indigenous Hoabinhian hunter-gatherers and Neolithic farmers who migrated southward from China (*1, 6, 29*).

### The 1000 Genomes Reference Panel

To broaden our analyses to additional Asian populations, we incorporated Asian populations from the 1000 Genomes Project Phase 3 (*31*). We downloaded the data in the hg38 reference build to ensure high coverage, but, since our data and the Neanderthal and Denisovan genomes are aligned to hg37, we converted the 1000 Genomes data to hg37 using Picard’s *LiftoverVcf* (*63*)

We then removed duplicated and monomorphic sites, as well as indels and non-biallelic sites using bcftools (*64*). Afterward, we removed the related individuals using KING (*65*). Finally, we merged this dataset with the archaic genomes of the Altai Neanderthal (*15*), Vindija Neanderthal (*11*), and Altai Denisova (Denisova 3 (*8, 30*)), as well as the chimpanzee reference genome (PanTro4) (*66*).

### Phasing and Imputation

To prevent ascertainment bias, we imputed genotype data after evaluating the impact of the imputation process on the level of Neanderthal ancestry (Supplementary text 1). We phased the data following the protocol of SHAPEIT4 (*64*), with the 1000 Genomes Project dataset excluding African populations as the reference panel. Similarly, SNP-array data were imputed following the protocol developed for Beagle5.4 (*65*), using the same reference panel as for phasing.

### Final datasets and quality control

We phased and imputed SNP-array data using the 1000 Genomes panel excluding African populations, yielding 1,005 whole-genome-equivalent individuals (referred to as the *modern dataset*). We merged this dataset with archaic genomes to build the *archaic dataset*. We removed sites in CpG islands (UCSC Table Browser) and segmental duplications.

For the archaic dataset, we additionally removed sites outside archaic accessibility masks (Vindija, Altai Neanderthal, Altai Denisovan; http://ftp.eva.mpg.de/neandertal/Vindija/FilterBed). This led to two datasets: the modern dataset and the archaic dataset, containing 54,165,660 SNPs and 25,244,313 SNPs, respectively.

### Estimating archaic ancestry

We computed D-statistics of the form *D(Chimpanzee, Vindija; YRI, X)* and *D(Chimpanzee, Denisova 3; CEU, X)* to assess sharing of derived alleles. Genome-wide ancestry proportions were estimated via f4-ratio:

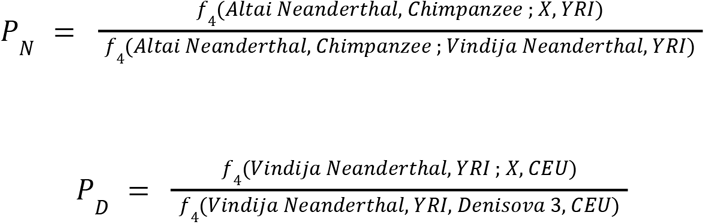

### Identifying introgressed haplotypes

#### Sprime

We applied Sprime (*4*) to detect segments enriched for variants rare or absent in Yoruba. Following Browning et al., we retained only segments with at least ten such SNPs and a detection score threshold of 150,000. We computed match rates to the Vindija Neanderthal and Altai Denisovan genomes using *maparch* (*68*), masking low-mappability/coverage regions and retaining segments with less than 30 unmasked sites. Segments were classified as Neanderthal if the Vindija match rate exceeded 0.6 and the Denisovan match rate was below 0.4; segments were classified as Denisovan if the Denisovan match rate exceeded 0.3 and the Vindija match rate was below 0.3.

#### CRF

To confirm these observations, we used a conditional random field (CRF) approach developed by Sankararaman et al. (*10*) that detects archaic introgressed haplotypes using an archaic reference genome. To identify Neanderthal introgressed haplotypes we used the African genomes from the 1000 Genomes Project (*31*) and the Denisova 3 genome (*8*) as outgroups. Conversely, to detect Denisovan introgressed haplotypes we used the African genomes from the 1000 Genomes Project (*31*) and the Vindija Neanderthal genome (*11*) as outgroups. A site was classified as Neanderthal-derived if its Neanderthal posterior probability was >0.9 and Denisovan posterior probability <0.5, and vice versa for Denisovan-derived sites. We retained segments containing at least 100 sites, and computed match rates as the proportion of sites matching either the Neanderthal or Denisova genome relative to the total number of sites in each segment. We kept segments with a match rate higher than 0.953, as the bimodal distribution already reported in East Asian populations was apparent and stable with this threshold, while using higher thresholds substantially reduced the number of segments available for analysis. To reduce false positives (*12, 16*), we only analysed segments jointly identified by Sprime and CRF. Finally, we fitted Gaussian mixture models (1< k < 10) to match-rate distributions.

### Dating Denisovan admixture event

We assigned haplotypes to admixture pulses and compared inter-population overlap and haplotype lengths using Wilcoxon rank-sum tests.

### Adaptive introgression

We considered only segments identified as introgressed by both Sprime and the CRF approach (11,285 Neanderthal segments and 370 Denisovan segments). We intersected their genomic coordinates with 5,566 regions previously identified as targets of positive selection using iHS (see (*69*)) using Bedtools *--multiinter (70)*. This resulted in 2,904 and 122 regions overlapping regions for Neanderthal and Denisovan segments, respectively. We then annotated these regions and retained only those spanning genes included in a curated list of 76 candidates previously reported as cases of adaptive introgression in the literature (*4, 12, 26, 52*).

We are grateful to the sampled populations for collaborating on this research. We thank Stéphane Peyrègne, Etienne Patin, Guillaume Laval, Isabelle Crèvecoeur and Marie-Claude Marsolier for useful comments and discussions. AI tools were used to rephrase certain sentences.

## Funding

ANR NUTGENEVOL (07-BLAN-0064); ANR Altérité culturelle (10-ESVS-0010)

Emergence programs HOMOCULTURGEN from Sorbonne Universités & Sorbonne Paris

Cité (SU-15-R-EMR-02) to E.H

ANR SoGen (JC09_441218) grant to RC

C. A-D and J. AD were founded by Ecole Doctorale 227, DIVONA

## Author contributions

Each author’s contribution(s) to the paper should be listed (we suggest following the CRediT model with each CRediT role given its own line. No punctuation in the initials.

Conceptualization: FD, BT, CB, EH, RC, CAD

Methodology: CAD, RL

Investigation: CAD

Supervision: RL, FD, BT, CB

Writing—original draft: CAD

Writing—review & editing: CAD, JAD, RL, FD, BT, CB, RC, EH

## Data and materials availability

The data for populations from CNA and SEA presented here can be accessed and downloaded via the European Genome-Phenome Archive (EGA) database under accession numbers EGAD00010002663, EGAD00010001724 and EGAD00010001538. The data can be shared provided that the future studies comply with the participants informed consents.

