## Supplementary texts and figures for "The genetic legacy of archaic hominins in Central and Southeast Asia uncovers three distinct Denisovan populations"

Charlotte Antoine-Derouet et al.

**This PDF file includes:**

Supplementary Text 1 to 5
Figs. S1 to S7
Tables S1 to S7

**Supplementary text 1**
**Testing phasing and imputation**
*Reference panels*
To mitigate SNP-array ascertainment bias, we performed genotype imputation and determined which was the best reference panel to use for imputation. We tested six different reference panels:

- Full 1000 Genome Reference Panel
- Europeans and East Asians from the 1000 Genome Reference Panel
- Whole Genome Sequencing data from the CTA, LPO, SHO, TJE and TUR
- 1000 Genome Reference Panel without Africans
- 1000 Genome Reference Panel without Africans + Whole Genome Sequencing data from LPO, SHO, TUR
- Full 1000 Genome Reference Panel + Whole Genome Sequencing data from LPO, SHO, TUR

We extracted SNPs at Omni1 positions from WGS individuals (CTA, LPO, TJE, TUR and SHO), harmonised alleles to the 1000 Genomes reference, removed allele mismatches, phased the data with SHAPEIT4 (*67*), and performed genotype imputation with Beagle5.4 (*68*) separately using each of the six reference panels.

*Evaluation*
For each panel, we merged the imputed datasets with our working dataset (1000 Genomes + archaic hominins + chimpanzee), retained overlapping positions, and generated ten random subsets of five million sites (70 datasets in total, 6 datasets for each imputation reference panel and a dataset composed of the Whole genome data). We estimated Neanderthal ancestry with f4-ratio statistic (see below) and compared results from imputed and non-imputed data. Differences were generally small; the lowest discrepancies were observed when imputing using 1000 Genomes Reference Panel without Africans or when whole-genome sequencing (WGS) data were included in the reference dataset (Fig. S1). However, to avoid potential bias, we excluded our WGS individuals from the final reference panel and used the 1000 Genomes dataset without Africans.

**Supplementary Text 2**
**Comparison of Sprime results using imputed and non-imputed data**
To further assess whether imputation affected our results, we ran Sprime three times on the CTA population, for which both WGS and SNP-array data are available. First, we applied Sprime on the full CTA dataset, comprising a mixture of individuals with non-imputed WGS data and imputed SNP-array data, and observed a three-pulse pattern of Denisovan ancestry. To evaluate whether this pattern could result from imputation, we repeated the analysis

using only non-imputed WGS individuals, and, separately, on imputed SNP-array data. In both cases, the three-pulse pattern was recovered, indicating that imputation does not affect our results (Fig. S2).

**Supplementary Text 3**

**Distribution of combined Sprime and CRF Denisovan introgressed haplotypes**

Before merging Denisovan haplotypes by region, we fitted Gaussian Mixture Models to the distribution of match rates to assess whether a three-pulse pattern could be detected despite the limited number of Denisovan introgressed haplotypes in Southeast Asian populations. We found that, in severalSoutheast Asian populations (CBR, CTA, LPO), a three-component model provides the  best fit to  the data. This supports the hypothesis that Denisovan ancestry in Southeast Asia results from three different admixture events involving three genetically differentiated Denisovan populations (Fig. S4).

**Supplementary Text 4**

**Distribution of Denisovan haplotypes among population from a same region**

We analysed the distribution of haplotypes assigned to different admixture pulses across populations to assess whether these signals were consistently represented or whether some haplotypes assigned to a specific pulse were detected in only a single population within a given region. Overall, we found that haplotypes are broadly shared across populations and that what we consider as a pulse is not haplotypes with a specific match rate in a single population but results from haplotypes identified in several populations from the same region (Fig. S5).

**Supplementary Text 5**

**Correlation between Sprime match rates and CRF match rates, length and number of sites**

To investigate whether the third peak reflects an additional admixture event with another Denisovan population, we tested whether match rate correlates with segment length. A significant correlation could indicate that shorter fragments systematically exhibit lower match rates (or vice versa), suggesting that the observed third pulse may result from fragment length-dependent biases rather than an additional Denisovan admixture event. We found no correlation between segment length and match rate in any population. In Southeast and East Asian populations, match rate showed a weak correlation with the number of sites per segment (p-value = $5.84 \times 10^{-5}$, p-value = 0.0097, respectively), although the effect size was small (r = 0.175, r = 0.154, respectively) (Fig. S6).

In addition, we assessed the consistency of segment identification across methods by comparing match rates computed by Sprime and CRF. We observed a significant correlation between the two methods in  East and Southeast Asian populations, indicating that segments identified by Sprime are largely consistent with those detected using CRF (Fig. S6).

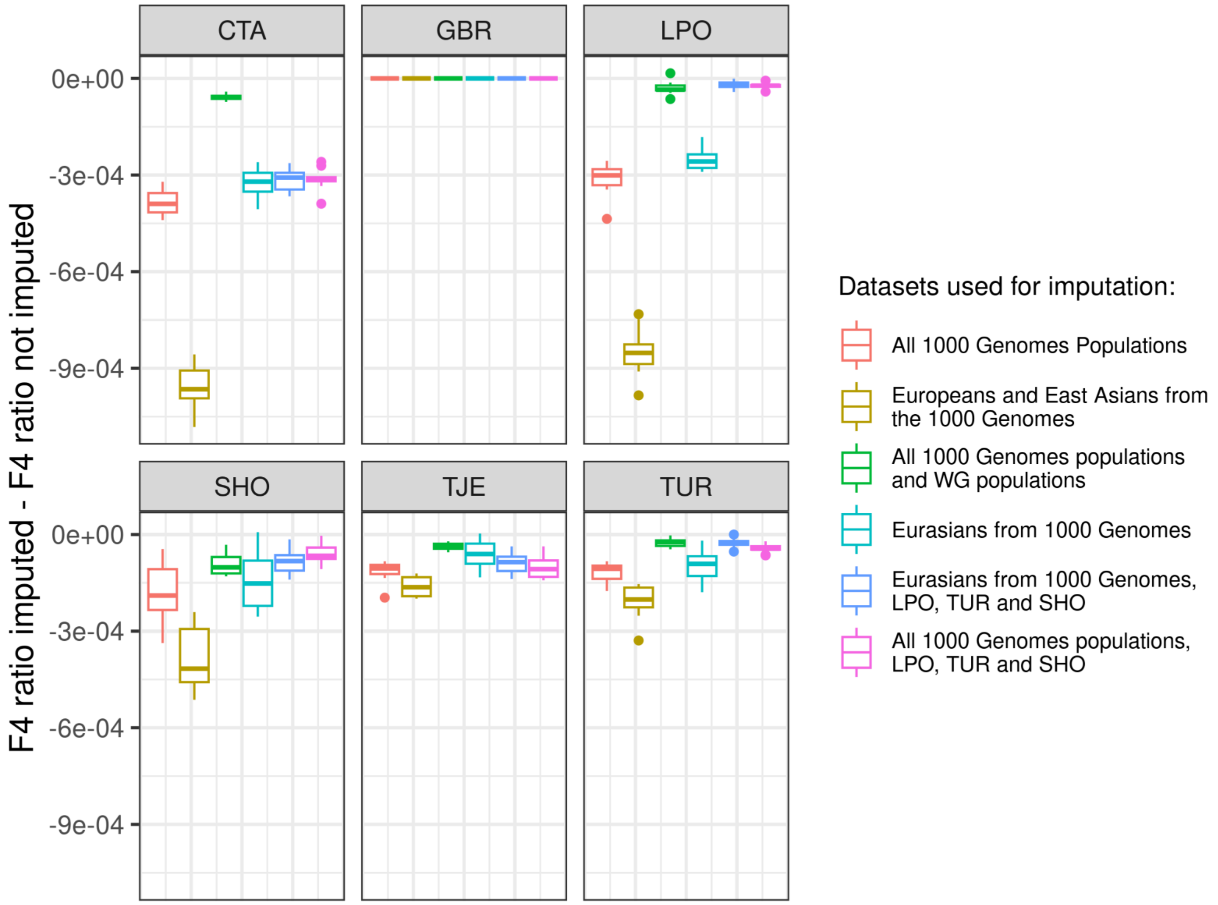


**Fig. S1. Boxplots showing differences between Neanderthal ancestry estimates between imputed and non-imputed datasets.**
Colors correspond to the reference panels used for phasing and imputation.

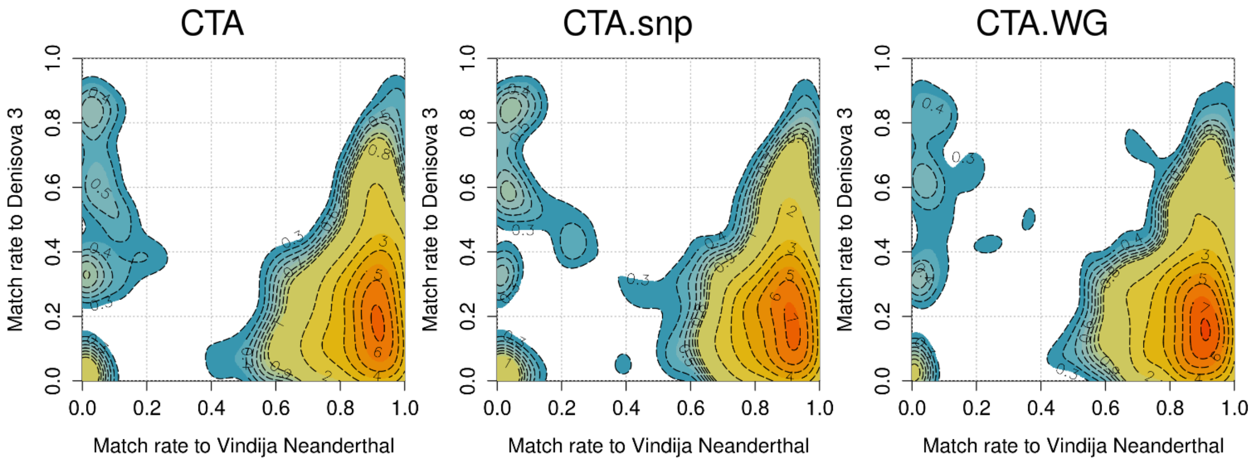


**Fig. S2. Contour density plots of Match proportion of introgressed segments to the Vindija Neanderthal and Denisova3 genome.**
CTA corresponds to results on combined CTA that were imputed and non-imputed. CTA.snp are the results of only imputed individuals. CTA.WG contains only WG individuals. The match rate is the proportion of

putative archaic alleles that match a given archaic genome, excluding sites at masked positions. Only Sprime haplotypes with at least 30 sites not masked in the Vindija Neanderthal and Denisova3 genomes are included in the match rate calculations. Numbers inside the contour plots indicate the height of the density corresponding to each contour line. Contour lines are shown for multiples of 1 (solid lines) and multiples of 0.1 between 0.3 and 0.9 (dashed lines). Colors were added to enhance the comprehension.

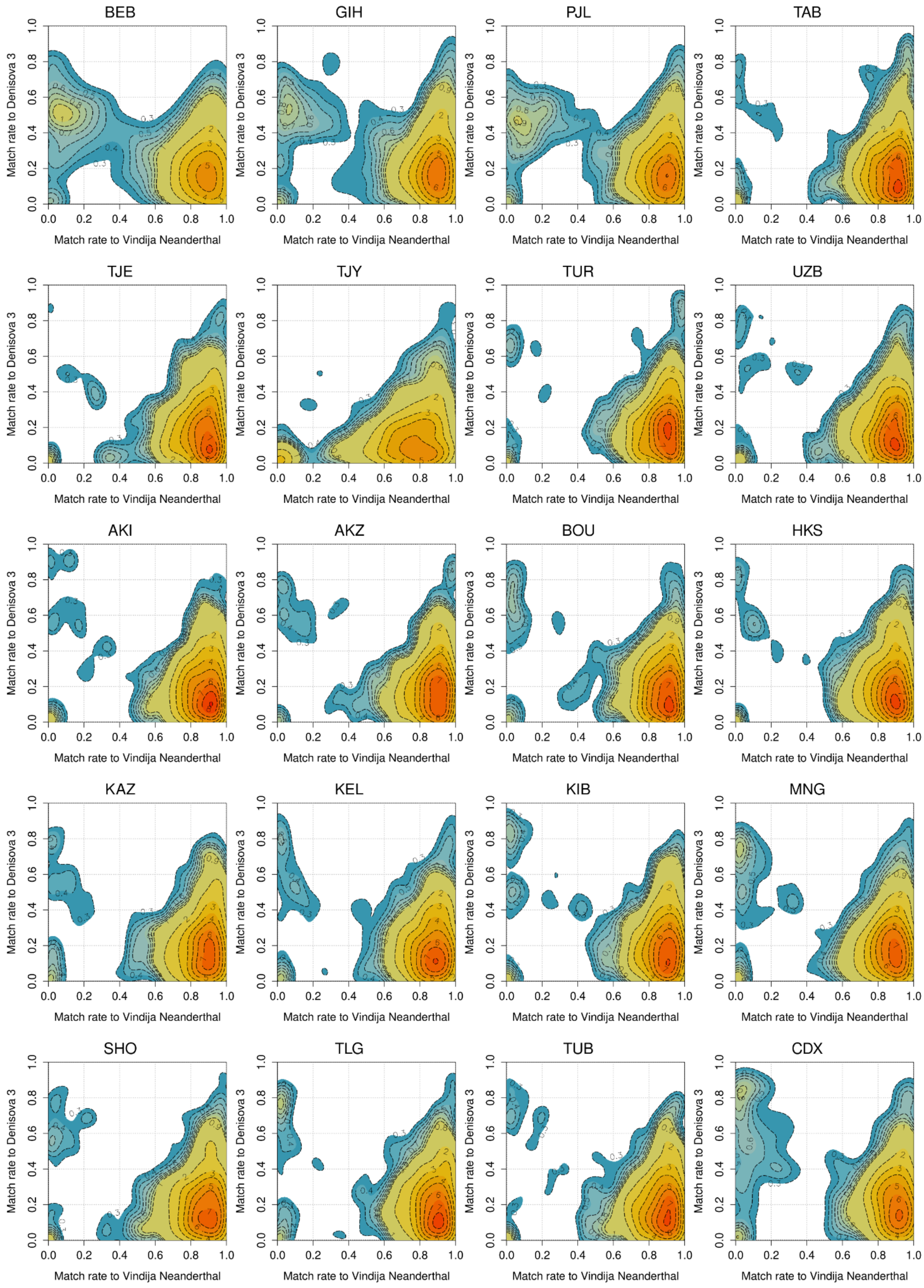
BEB
GIH
PJL
TAB
TJE
TJY
TUR
UZB
AKI
AKZ
BOU
HKS
KAZ
KEL
KIB
MNG
SHO
TLG
TUB
CDX
Match rate to Denisova 3
Match rate to Vindija Neanderthal
0.0
0.2
0.4
0.6
0.8
1.0

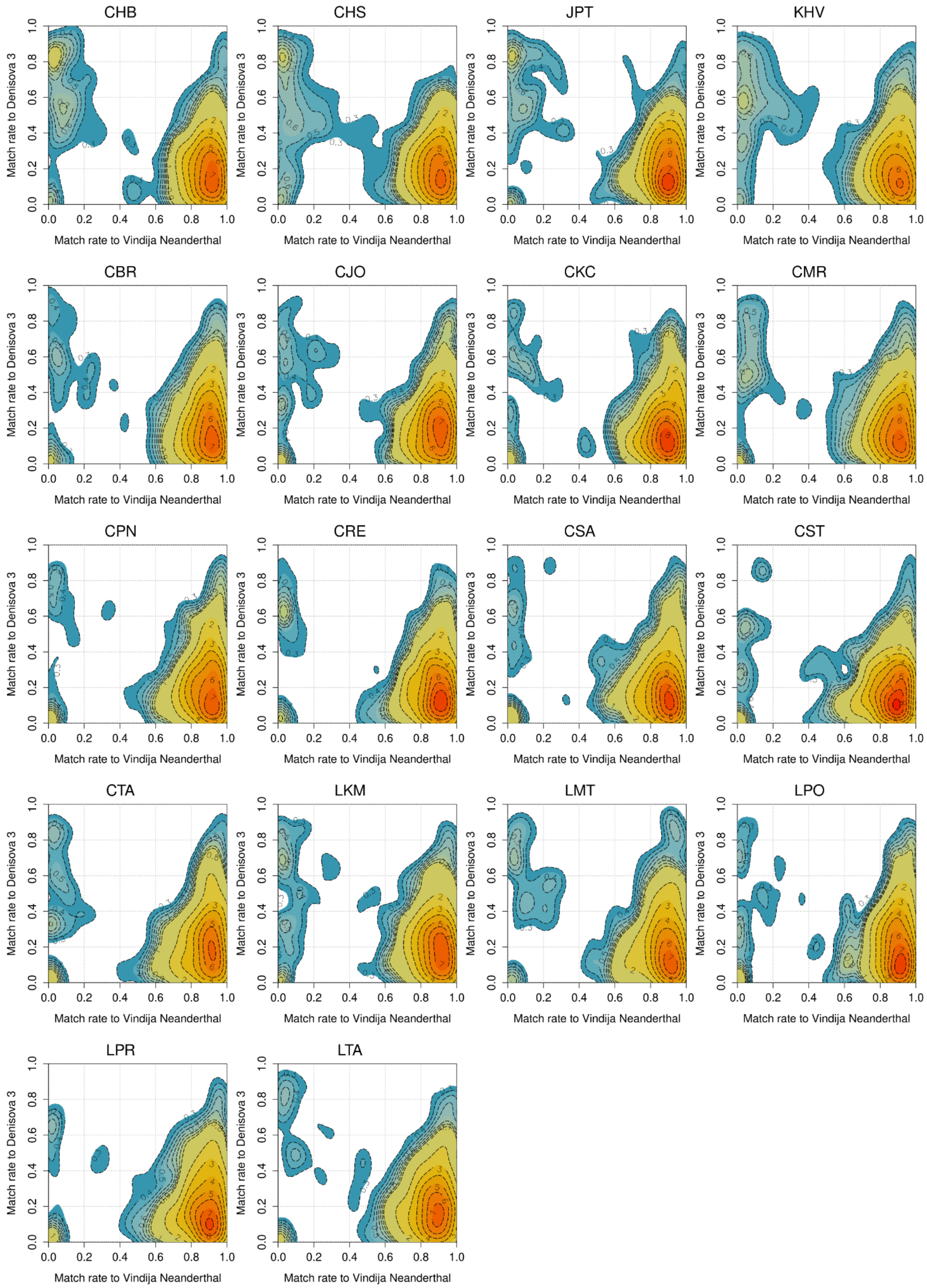
CHB
CHS
JPT
KHV
CBR
CJO
CKC
CMR
CPN
CRE
CSA
CST
CTA
LKM
LMT
LPO
LPR
LTA
Match rate to Denisova 3
Match rate to Vindija Neanderthal
0.0
0.2
0.4
0.6
0.8
1.0

**Fig. S3. Contour density plots of Match proportion of introgressed segments to the Vindija Neanderthal and Denisova3 genome.**

The match rate is the proportion of putative archaic alleles that match a given archaic genome, excluding sites at masked positions. Only Sprime haplotypes with at least 30 sites not masked in the Vindija Neanderthal and Denisova3 genomes are included in the match rate calculations. Numbers inside the contour plots indicate the height of the density corresponding to each contour line. Contour lines are shown for multiples of 1 (solid lines) and multiples of 0.1 between 0.3 and 0.9 (dashed lines). Color were added to enhance the comprehension

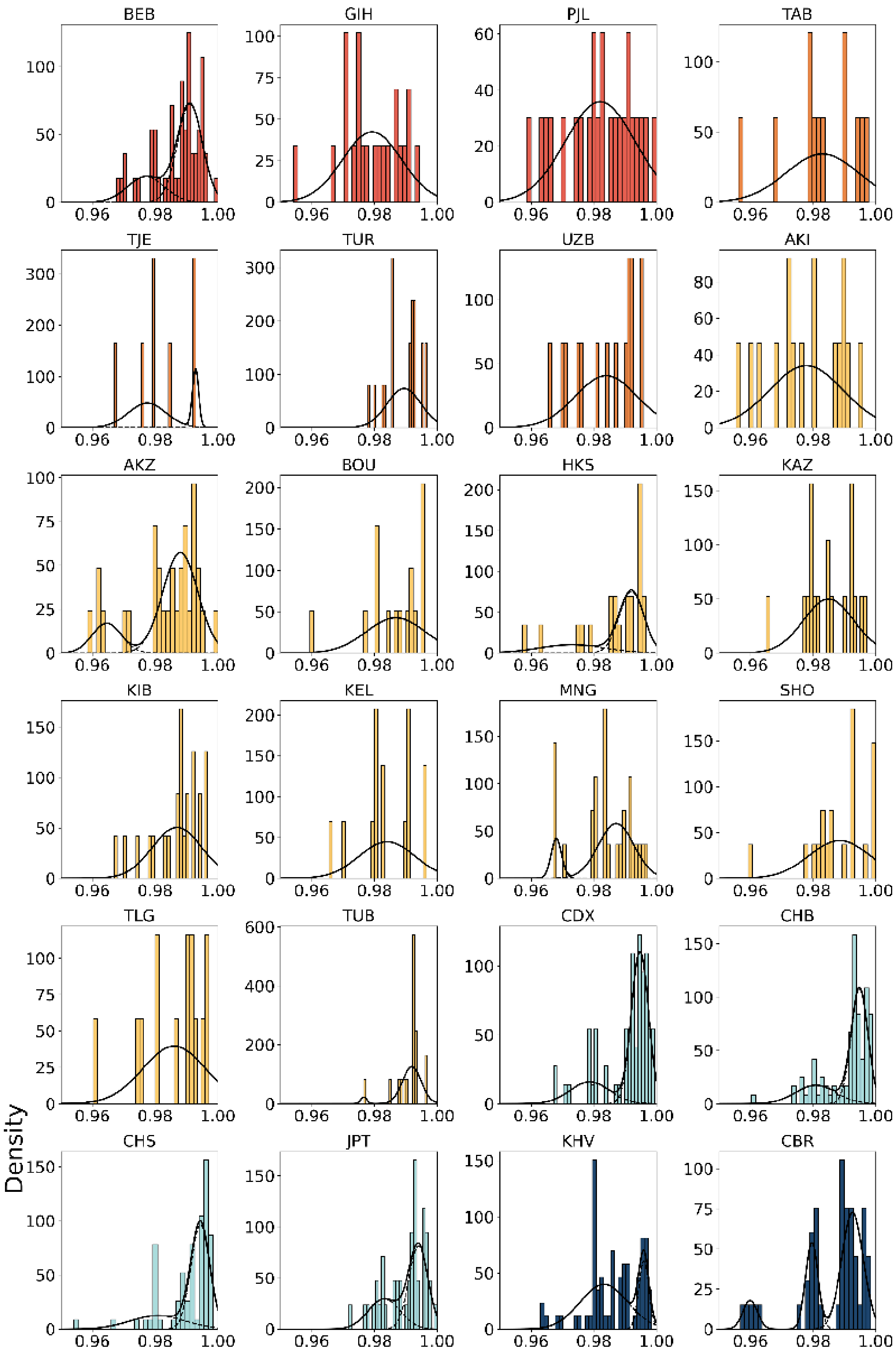

BEB
GIH
PJL
TAB
TJE
TUR
UZB
AKI
AKZ
BOU
HKS
KAZ
KIB
KEL
MNG
SHO
TLG
TUB
CDX
CHB
CHS
JPT
KHV
CBR
Density
0.96
0.98
1.00

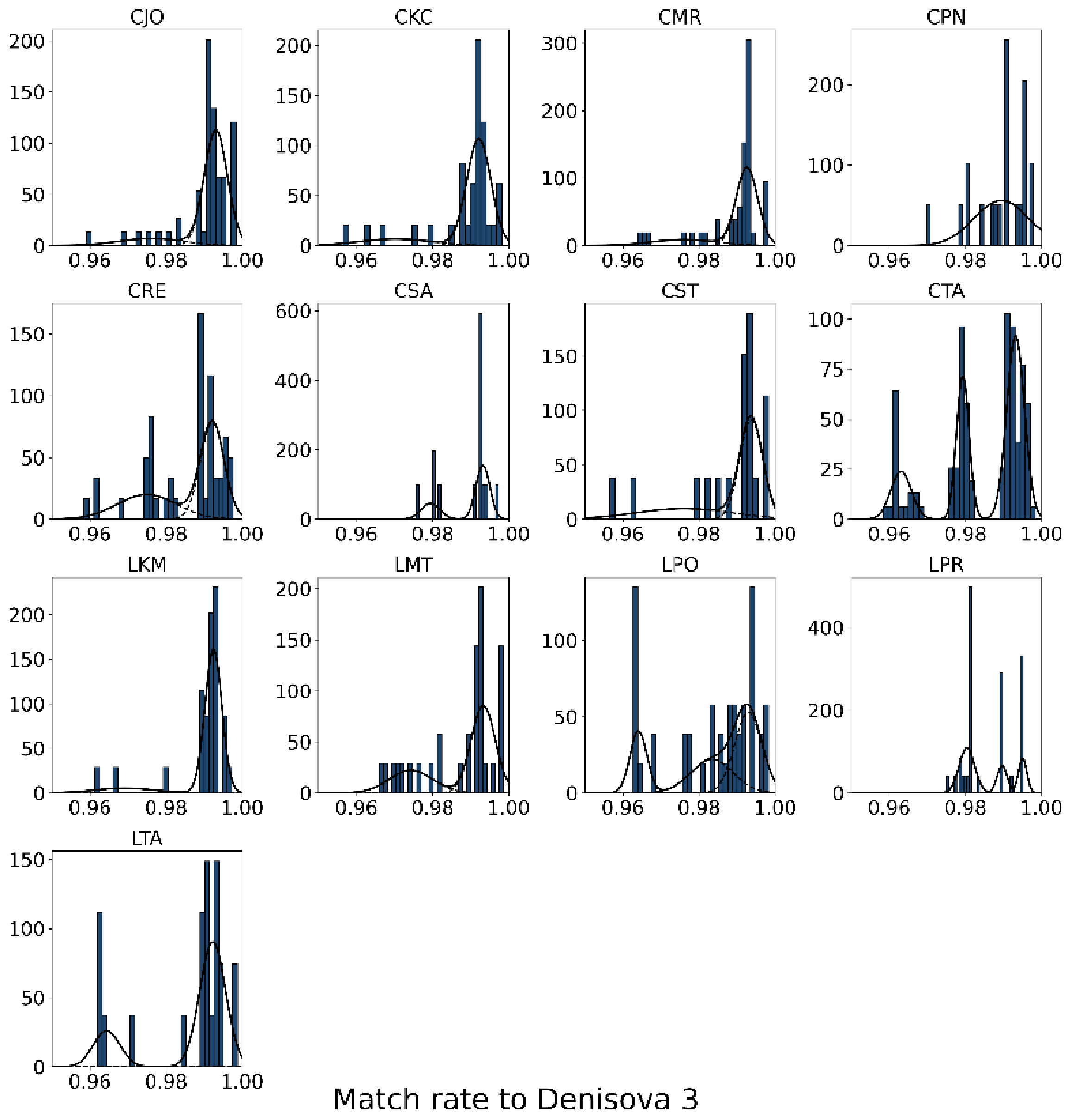


**Fig. S4. Match rate distributions of Denisovan introgressed haplotypes.**
Here we plotted and fit Gaussian mixture on haplotypes that are identified by Sprime and CRF. Colors correspond to the genetic geographic attribution.

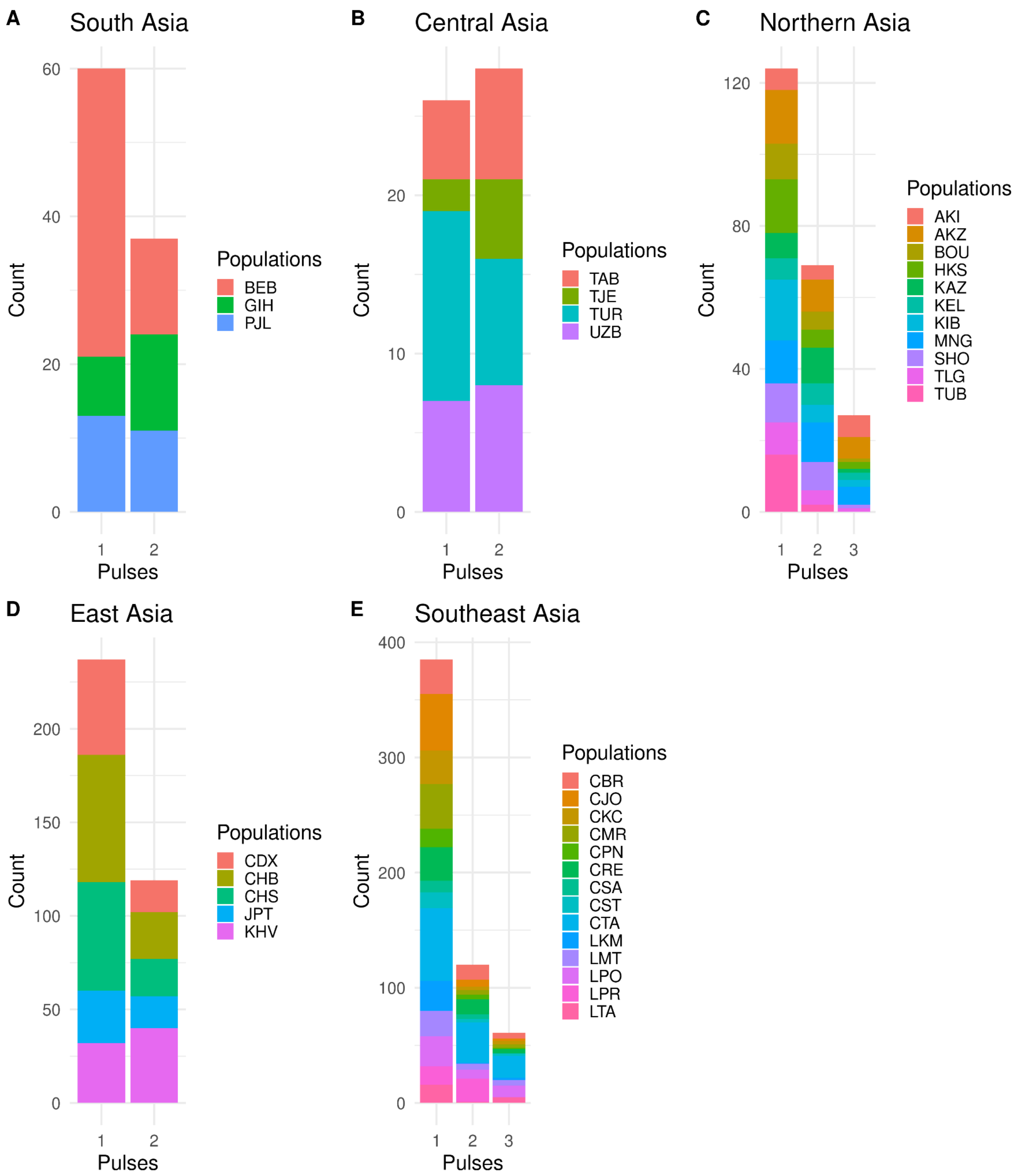


**Fig. S5. Distribution of the number of haplotypes assigned to each admixture pulse across regions.**
The pulses are numbered in descending order of match rate

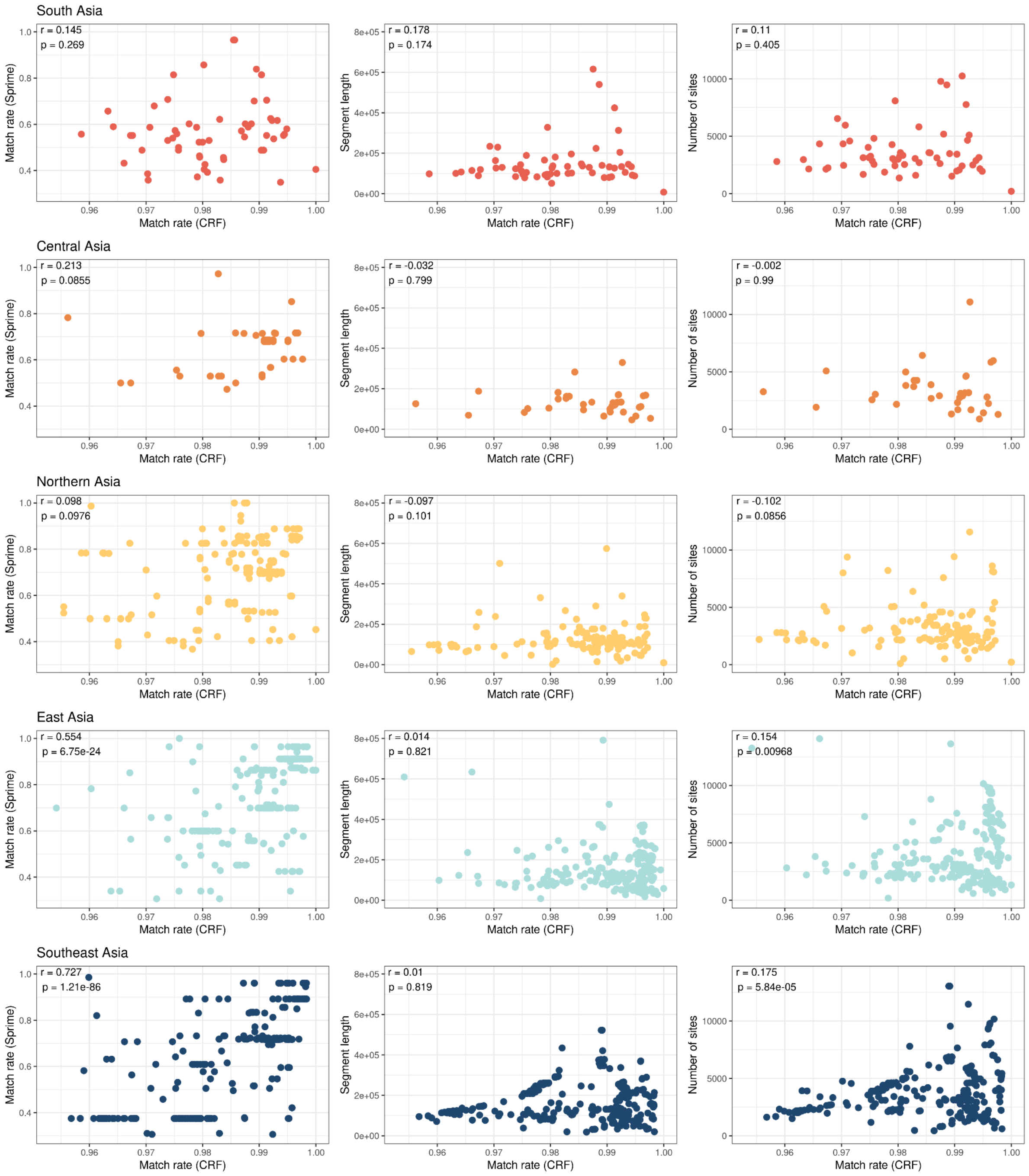


**Fig. S6. Correlation analyses.**
Left: Comparison of match rates estimated by CRF and Sprime. Middle: Relationship between match rates estimated by CRF and haplotype lengths. Right: Relationship between match rates estimated by CRF and the number of sites per segment.

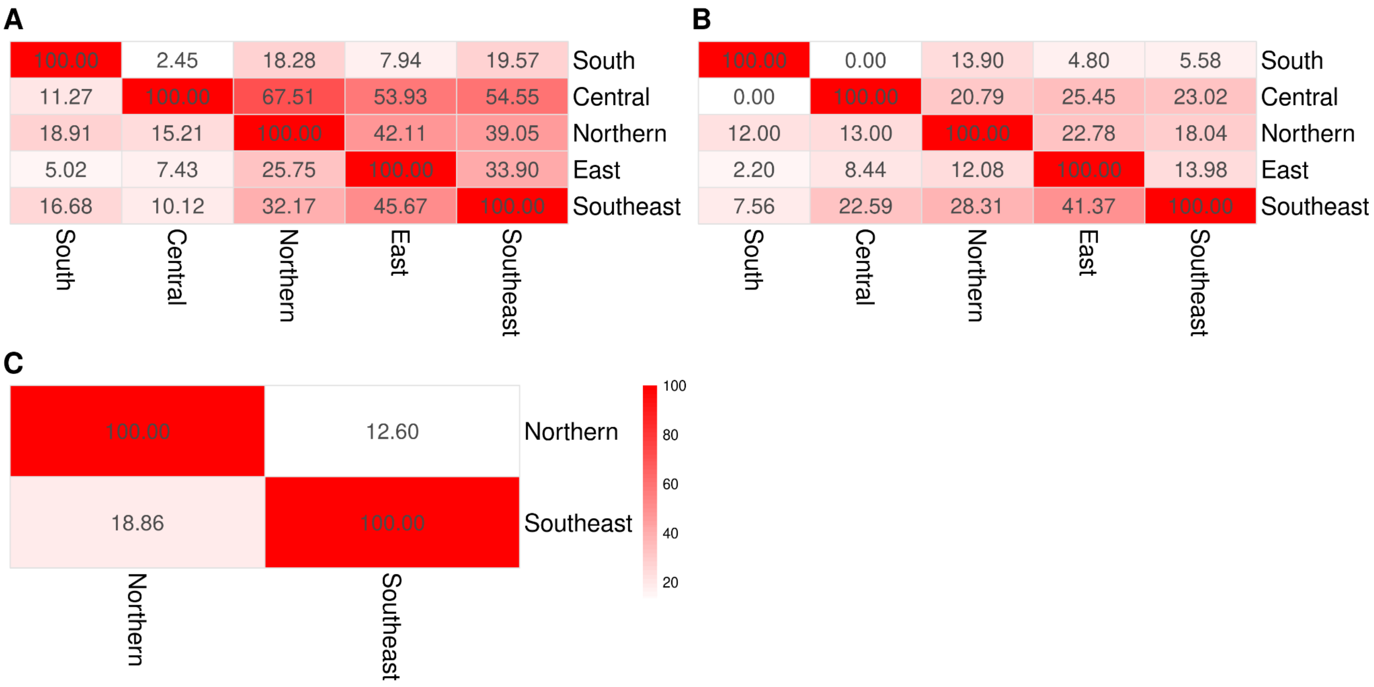


**Fig. S7. Matrix showing haplotype overlap between regions.**
**A**. Pulse1 (highest match rate). **B**. Pulse2 (moderate match rate). **C**. Pulse3 (lowest match rate). Overlap is computed as the number of bases in column region that overlap row region divided by the number of bases in the row region, multiplied by 100.
